# Delimiting Continuity: Comparison of Target Enrichment and ddRAD for Delineating Admixing Parapatric *Melitaea* Butterflies

**DOI:** 10.1101/2022.02.05.479083

**Authors:** Mukta Joshi, Marianne Espeland, Vlad Dincă, Roger Vila, Mohadeseh S. Tahami, Lars Dietz, Christoph Mayer, Sebastian Martin, Leonardo Dapporto, Marko Mutanen

## Abstract

Parapatrically distributed taxa pose a challenge for species delimitation due to the presence of gene flow and inherent arbitrariness of exactly defining the species boundaries in such systems. We tackled the problem of species delimitation in a parapatric species pair of *Melitaea* butterflies using two popular genomic methods – double digest restriction-site associated DNA sequencing (ddRAD) and target enrichment. The former is mainly applied at shallow phylogenetic scales and the latter at both deep and shallow scales. Although both of these methods have been adequately utilised for species delimitation purposes, there are only a handful of studies that have compared these two genomic approaches in the same study system. We applied phylogenetic, population genetic and species delimitation methods and compared the results obtained from the two approaches. Using a recently developed target enrichment probe kit, we were able to capture 1,743 loci with a low amount of missing data and compared these with already available ddRAD data from a previous study on the same set of specimens. We recovered consistent phylogenetic relationships across the datasets, both demonstrating the presence of a genetically distinct Balkan lineage and paraphyly of *Melitaea athalia* with respect to *Melitaea celadussa.* The same relationships were also found in a species tree analysis of the target enrichment dataset using ASTRAL. Population genetic STRUCTURE analyses supported the presence of two species when using ddRAD data, but three species when using target enrichment, while the Bayes factor delimitation analysis found both two and three species scenarios equally decisive in both datasets. From the geographic distribution of genomic admixture, we confirm the patterns observed by a previous study that used ddRAD data. As the results obtained from both methods were largely congruent, we discuss some practical considerations and benefits of target enrichment over RAD sequencing. We conclude that the choice of method of genomic data collection does not influence the results of phylogenetic analyses at alpha taxonomic level, given a sufficient number of loci. Finally, we recommend a solution for delineating species in parapatric scenarios by proposing that parapatric taxa be consistently classified as subspecies or complete species, but not both, to promote taxonomic stability.

## INTRODUCTION

Most biologists would likely agree that, of all the taxonomic units, that of ‘species’ is fundamental (de Queiroz 2005; Hohenegger 2014). The reality of species as a truly discrete biological entity is however unlikely to hold true universally as speciation is usually a long and gradual process (de Queiroz 2007). The ability to categorize biological diversity into distinct species, i.e. species delimitation, is a prerequisite for the identification of individual organisms. The latter, in turn, is important in most biological research, including conservation, but also in many other disciplines and areas of society, such as legislation and food industry (Tewksbury et al. 2014). Despite the global efforts to document and describe the bewildering diversity of life on earth, most species remain undescribed or even undiscovered (Mora et al. 2011). In addition to the complexities caused by the usually low differentiation of recently diverged species, traits such as phenotypic plasticity and morphological crypticity displayed by some species seriously complicate the process of species delimitation (Knowlton 1993; Jarman and Elliott 2000). Delimitation is further complicated by several other factors. For instance, many species have a large or patchy distribution and show extensive intraspecific variability in geographical space. Additionally, hybridization and introgression between species have been shown to be much more common than previously expected. It has been estimated that 10% of animal species and at least 25% of all plant species are involved in hybridization (Mallet 2005). Finally, there is a variety of ways to define the term ‘species’. Some species concepts have a focus on temporal changes (i.e., when speciation occurs in time) while others focus on spatial (when geographic isolation between diverging populations) or biological (reproductive isolation) aspects of variability (de Queiroz 2007), rendering the definition of species even more elusive. Despite these difficulties, categorizing biodiversity into species remains a central commitment of taxonomy.

A wide geographical distribution or a geographically biased taxonomic sampling often complicates the process of species delimitation to a significant degree (Mutanen et al. 2012; Linck et al. 2019). When the distributions of species are large, extensive genetic differences among populations in different areas can be maintained, despite migration (Gavrilets et al. 2000). The same may be true on a smaller scale when dispersal is limited by species’ own traits or geographic barriers. In European butterflies, the degree of population differentiation is positively correlated with range size and negatively with traits determining their mobility and level of generalism (Dapporto et al. 2019). Also, variation in environmental conditions can result in selection acting differently in different areas within species’ range (Gavrilets et al. 2000). The geographic relationship between species or populations has traditionally been categorized as either allopatric, parapatric or sympatric (Hendry et al. 2009) although there is a full continuity between these distributional categories. Parapatry refers to the distributional pattern where the separate ranges of two species have a narrow overlap (Bull 1991), typically within a hybridisation zone (Hewitt 1988). While such circumstances provide an opportunity to peek into the speciation process through reinforcement, parapatry, in the presence of gene flow, poses a particular challenge for species delimitation, as the end point of the speciation process is often vague and gene flow may be extensive (Hendry et al. 2009). Indeed, the well-documented cases of parapatry demonstrate how utterly complicated species delimitation under such circumstances can be. A well-known example of such a system is that of carrion and hooded crows, *Corvus corone* (Linnaeus, 1758) and *Corvus cornix* (Linnaeus, 1758), in Europe (Saino and Villa 1992; Saino et al. 1992; Poelstra et al. 2013). These two taxa are now regarded as two different species (Saino et al. 1992) despite high levels of interbreeding and introgression.

Development of high-throughput sequencing technologies has enabled access to whole genomes or large proportions of genomes that can potentially provide useful information to resolve the patterns of evolution from population to very deep phylogenetic levels (Herrera and Shank 2016; Breinholt et al. 2018). Two of the popular reduced representation approaches, double digest restriction-site associated DNA sequencing (ddRAD; Peterson et al. 2012) and target capture methods such as ultraconserved elements (UCEs; Faircloth et al. 2012), anchored hybrid enrichment (AHE; Lemmon et al. 2012) and target enrichment (Gnirke et al. 2009, see also Mayer et al. 2016), are now widely used in systematics research. RADseq approaches have been used particularly to elucidate evolutionary patterns at relatively shallow phylogenetic levels (Wagner et al. 2013), while target capture-based approaches can be used for deep as well as shallow level phylogenomics (Banker et al. 2020).

The species pair *Melitaea athalia* (Rottemburg, 1775) and *Melitaea celadussa* (Fruhstorfer, 1910) represents a well-documented case of widespread and parapatric taxa in European butterflies with a hybrid zone mainly occurring in France, Switzerland, Italy and Austria, with *M. celadussa* occurring south and west, and *M. athalia* north and east of the contact zone (Higgins 1955; Coutsis & Van Oorschot 2014; Platania et al. 2020). The species are morphologically distinguished based on male genitalia, but intermediate male genital characters have been reported from the contact zone (Higgins 1955; Oorschot & Coutsis 2014). Genomic analyses done using a ddRAD dataset on this species pair revealed wide admixture in the contact zone and across a part of the range of *M. athalia*, but less so in *M. celadussa*, suggesting a mostly unidirectional introgression (Tahami et al. 2021). Additionally, this analysis revealed a genetically diverging and non-admixed, but morphologically indistinguishable, clade in the Balkans that had not been detected previously. Using morphometrics of the male genitalia, the same study reported several specimens with intermediate genital characters in the contact zone.

As parapatric systems are likely to reveal a full spectrum of genetic admixture across different systems, taxonomic delimitation of parapatric taxa is bound to be almost inherently arbitrary. However, despite the conceptual nature of their delimitation, there is no consensus in the taxonomic community on the principles of how such taxa should be taxonomically ranked. We believe that genomic approaches could provide excellent means for higher levels of standardization for delimiting parapatric taxa by enabling the evaluation of important biological properties, such as levels of introgression and genetic divergence, in a quantitative way and based on large amounts of standardized genetic markers (Dietz et al. 2019; Eberle et al. 2020). Hence, motivated by the low number of studies that use genomics to assess parapatric species systems, we tackled the question of their delimitation based on two powerful genomic approaches, ddRAD sequencing and target enrichment. We compared the two methods making use of phylogenetic, population genetic and species delimitation methods, because species delimitation essentially acts at the interface of phylogenetic and population genetic levels. Apart from comparison of the results and usefulness of ddRAD sequencing vs. target enrichment, we provide insights into the conceptual aspects of the delimitation of parapatric taxa with admixture in order to advance the stability and consistency of their delimitation.

## METHODS

### Sample preparation

Samples were collected from various locations across Europe and were representative for the range of the targeted taxa, including their contact zone (Table 1). Initially, a set of 81 specimens was used for ddRAD analyses (Tahami et al. 2021) and a subset of 60 specimens was selected for target enrichment analyses. These included specimens of *M. athalia* and *M. celadussa*, as well as one specimen putatively attributed to *Melitaea caucasogenita* (Verity, 1930) and one *Melitaea britomartis* (Assmann, 1847) as outgroup taxa. Because some specimens were females, it was not clear how well they can be identified based on genitalia, DNA barcodes were used for a-priori identification of specimens for communication purposes. We also categorized specimens occurring in the supposed contact zone using the biodecrypt approach (Platania et al. 2020), which implements dedicated functions to identify areas of sympatry among cryptic taxa based on a subset of identified specimens (details are given in the supplementary file). As per this analysis, we inferred 11 specimens as belonging to the contact zone (supplementary table S2). Genomic DNA was extracted from two thirds of the thorax from either ethanol preserved or dry specimens. DNA extraction was done using the QIAGEN DNeasy Blood and Tissue kit following the protocol provided by the manufacturer. DNA extracts were visualized on 1% agarose gels.

**Table 1:** Sample overview and comparison of number of raw reads from the target enrichment and ddRAD datasets

| Specimen ID | BioSample accessions | Taxon based on COI | Country | No. of target enrichment reads after filtering | No. of RAD reads after filtering |
| --- | --- | --- | --- | --- | --- |
| **RVcoll07E394** | SAMN25488010 | *Melitaea athalia* | Romania | 2072173 | 4673359 |
| **RVcoll08M346** | SAMN25488011 | *Melitaea athalia* | Romania | 2001739 | 1180010 |
| **RVcoll10A789** | SAMN2548801<br>2 | *Melitaea*<br>*athalia* | Estonia | 2219329 | 136195<br>8 |
| **RVcoll12Z197** | SAMN2548801<br>3 | *Melitaea*<br>*athalia* | Sweden | 1991064 | 342852<br>0 |
| **RVcoll13U124** | SAMN2548801<br>4 | *Melitaea*<br>*celadussa* | Italy | 2050917 | 104716<br>2 |
| **RVcoll13U296** | SAMN2548801<br>5 | *Melitaea*<br>*athalia* | Italy | 1836742 | 234894<br>4 |
| **RVcoll13U438** | SAMN2548801<br>6 | *Melitaea*<br>*athalia* | Italy | 2084530 | 179393<br>9 |
| **RVcoll14B773** | SAMN2548801<br>7 | *Melitaea*<br>*athalia* | Albania | 1614654 | 133406<br>2 |
| **RVcoll14D059** | SAMN2548801<br>8 | *Melitaea*<br>*athalia* | Bulgaria | 1720383 | 290442<br>2 |
| **RVcoll14E853** | SAMN2548801<br>9 | *Melitaea*<br>*athalia* | Serbia | 1396629 | 173521<br>4 |
| **RVcoll14E859** | SAMN2548802<br>0 | *Melitaea*<br>*athalia* | Serbia | 1605420 | 133643<br>9 |
| **RVcoll14E904** | SAMN2548802<br>1 | *Melitaea*<br>*athalia* | Serbia | 1703860 | 312305 |
| **RVcoll14F303** | SAMN2548802<br>2 | *Melitaea*<br>*athalia* | Serbia | 1885222 | 514133 |
| **RVcoll14F407** | SAMN2548802<br>3 | *Melitaea*<br>*athalia* | Bulgaria | 1464925 | 526291 |
| **RVcoll14F538** | SAMN2548802<br>4 | *Melitaea*<br>*athalia* | Greece | 1827776 | 141875<br>2 |
| **RVcoll14F650** | SAMN2548802<br>5 | *Melitaea*<br>*athalia* | Greece | 1470771 | 800882 |
| **RVcoll14F666** | SAMN2548802<br>6 | *Melitaea*<br>*athalia* | Greece | 1710058 | 899141 |
| **RVcoll14G434** | SAMN2548802<br>7 | *Melitaea*<br>*athalia* | Greece | 1830124 | 822392 |
| **RVcoll14V075** | SAMN2548802<br>8 | *Melitaea*<br>*athalia* | Ukraine | 1819607 | 263974<br>3 |
| **RVcoll15I360** | SAMN2548802<br>9 | *Melitaea*<br>*athalia* | Austria | 1736020 | 344731<br>4 |
| **RVcoll15P033** | SAMN2548803<br>0 | *Melitaea*<br>*athalia* | Ukraine | 1945474 | 239457<br>3 |
| **RVcoll16H415** | SAMN2548803<br>1 | *Melitaea*<br>*athalia* | Sweden | 1757794 | 663042<br>6 |
| **RVcoll16I052** | SAMN2548803<br>2 | *Melitaea*<br>*athalia* | Poland | 1624217 | 139973<br>8 |
| **RVcoll16J000** | SAMN2548803<br>3 | *Melitaea*<br>*athalia* | Slovakia | 1821930 | 166926<br>9 |
| **RVcoll16J612** | SAMN2548803<br>4 | *Melitaea*<br>*athalia* | Russia | 1927534 | 347774<br>8 |
| **MAT-SG-W-**<br>**144** | SAMN2548803<br>5 | *Melitaea*<br>*celadussa* | Switzerlan<br>d | 1721973 | 220727<br>4 |
| **RVcoll08J851** | SAMN2548803<br>6 | *Melitaea*<br>*celadussa* | Spain | 1726868 | 213919<br>1 |
| **RVcoll08L852** | SAMN2548803<br>7 | *Melitaea*<br>*celadussa* | Spain | 1538438 | 116216<br>1 |
| **RVcoll08M07**<br>**4** | SAMN2548803<br>8 | *Melitaea*<br>*celadussa* | Spain | 1880216 | 123379<br>8 |
| **RVcoll08M91**<br>**5** | SAMN2548803<br>9 | *Melitaea*<br>*celadussa* | Spain | 1735217 | 131998<br>7 |
| **RVcoll08P221** | SAMN2548804<br>0 | *Melitaea*<br>*celadussa* | Spain | 1942107 | 236183<br>5 |
| **RVcoll11H561** | SAMN2548804<br>1 | *Melitaea*<br>*celadussa* | Italy | 1676314 | 912613 |
| **RVcoll11H741** | SAMN2548804<br>2 | *Melitaea*<br>*celadussa* | Italy | 1617340 | 212797<br>4 |
| **RVcoll11I507** | SAMN2548804<br>3 | *Melitaea*<br>*celadussa* | Spain | 1835635 | 168284<br>8 |
| **RVcoll11I949** | SAMN2548804<br>4 | *Melitaea*<br>*celadussa* | France | 1843398 | 108721<br>5 |
| **RVcoll12O623** | SAMN2548804<br>5 | *Melitaea*<br>*celadussa* | France | 1365257 | 184401<br>4 |
| **RVcoll12P926** | SAMN2548804<br>6 | *Melitaea*<br>*celadussa* | France | 2152092 | 180775<br>7 |
| **RVcoll12Q105** | SAMN2548804<br>7 | *Melitaea*<br>*celadussa* | France | 1799934 | 190377<br>1 |
| **RVcoll12Q106** | SAMN2548804<br>8 | *Melitaea*<br>*celadussa* | France | 1818742 | 782461 |
| **RVcoll13S845** | SAMN2548804<br>9 | *Melitaea*<br>*celadussa* | Portugal | 1769707 | 417817<br>3 |
| **RVcoll13U092** | SAMN2548805<br>0 | *Melitaea*<br>*celadussa* | Italy | 2030995 | 236511<br>9 |
| **RVcoll14E220** | SAMN2548805<br>1 | *Melitaea*<br>*celadussa* | Spain | 1915377 | 505729<br>9 |
| **RVcoll14J820** | SAMN2548805<br>2 | *Melitaea*<br>*celadussa* | France | 1838442 | 291952<br>5 |
| **RVcoll14L240** | SAMN2548805<br>3 | *Melitaea*<br>*celadussa* | Italy | 1596947 | 229372<br>4 |
| **RVcoll15A654** | SAMN2548805<br>4 | *Melitaea*<br>*celadussa* | Italy | 2104662 | 684105 |
| **RVcoll15G145** | SAMN2548805<br>5 | *Melitaea*<br>*celadussa* | France | 1772903 | 109963<br>7 |
| **RVcoll15G841** | SAMN2548805<br>6 | *Melitaea*<br>*celadussa* | Italy | 1952921 | 971078 |
| **RVcoll15I495** | SAMN2548805<br>7 | *Melitaea*<br>*celadussa* | Austria | 1850398 | 136556<br>7 |
| **RVcoll15L146** | SAMN2548805<br>8 | *Melitaea*<br>*celadussa* | Italy | 1741727 | 273881 |
| **RVcoll15M13**<br>**3** | SAMN2548805<br>9 | *Melitaea*<br>*celadussa* | France | 1625035 | 296288<br>0 |
| **RVcoll15N014** | SAMN2548806<br>0 | *Melitaea*<br>*celadussa* | Italy | 2298510 | 291196<br>2 |
| **RVcoll16C754** | SAMN2548806<br>1 | *Melitaea*<br>*celadussa* | Italy | 1436688 | 818831 |
| **MAT-UR-I-**<br>**146** | SAMN2548806<br>2 | *Melitaea*<br>*celadussa* | Switzerlan<br>d | 1802507 | 148154<br>1 |
| **MAT-LU-K-**<br>**122** | SAMN2548806<br>3 | *Melitaea*<br>*celadussa* | Switzerlan<br>d | 1646303 | 151122<br>6 |
| **MAT-SG-W-**<br>**135** | SAMN2548806<br>4 | *Melitaea*<br>*celadussa* | Switzerlan<br>d | 1887704 | 163048<br>8 |
| **MAT-SG-W-**<br>**137** | SAMN2548806<br>5 | *Melitaea*<br>*celadussa* | Switzerlan<br>d | 1740049 | 863860 |
| **MAT-SG-W-138** | SAMN2548806<br>6 | *Melitaea*<br>*celadussa* | Switzerlan<br>d | 1742294 | 130880<br>9 |
| **MAT-SG-W-140** | SAMN2548806<br>7 | *Melitaea*<br>*celadussa* | Switzerlan<br>d | 1661150 | 489925 |
| **ZFMK-TIS-8000434** | SAMN2548806<br>8 | *Melitaea*<br>*caucasogenit*<br>*a* | Georgia | 1760707 | 203481<br>3 |
| **RVcoll14V076** | SAMN2548806<br>9 | *Melitaea*<br>*britomartis* | Ukraine | 1466749 | 195312<br>6 |

### Target enrichment laboratory procedures

Target enrichment bait design followed Mayer et al. (2021) where the final probe kit targets 2953 CDS regions in 1753 nuclear genes. This kit was developed with BaitFisher version 1.2.8 (Mayer et al. 2016) and is referred to as the LepZFMK 1.0 kit. The DNA concentration of each sample was quantified using a Promega Quantus fluorometer, and the initial fragment lengths were measured with a Fragment Analyzer (Agilent Technologies Inc.). About 100 ng absolute amount of DNA was taken for further processing as per standard Agilent protocol. The genomic DNA was subjected to random mechanical shearing to an average size between 250-300 bp followed by an end-repair reaction and ligation of adenine residue to the 3’ end of the blunt fragments (A-tailing) to allow ligation of barcoded adaptors using the Agilent SureSelect XT2 Library prep kit. Adaptor ligation and indexing was performed using the New England Biolabs reaction kit to enable dual indexed libraries. The resulting libraries were then PCR amplified. The number of PCR cycles at this stage was determined on the basis of library concentration after A-tailing and a total of 8 cycles were used for all samples. The concentration of PCR-amplified libraries was checked using the Quantus fluorometer and fragment distribution was analysed using the Fragment Analyzer. After the library construction, custom SureSelect baits (Agilent Technologies, SureSelect Custom Baits size 6-11.9 Mb) were used for solution-based target enrichment of a pool containing 8 libraries. The hybridisation was performed on each pool with SureSelect XT2 pre-capture ILM module following the manufacturer’s instructions. Enriched libraries were then captured with Streptavidin beads (MyOne Streptavidin T1). These captured libraries were PCR amplified and the final concentration of each captured library was again checked on the Quantus fluorometer. Following the enrichment, pooled libraries were sequenced using Illumina Nextseq 500 mid output to generate paired-end 150-bp reads at Starseq GmbH (Mainz, Germany).

### Target Enrichment bioinformatics

Adaptor sequences and low-quality regions were trimmed from demultiplexed data with fastq-mcf (Aronesty, 2011). The reads were mapped against the reference gene alignments of the bait regions generated during bait design with the BWA-MEM algorithm in bwa 0.7.17 (available from bio-bwa.sourceforge.net) with the minimum seed length set to 30. Reads for which the mapping was successful were then extracted from the resulting SAM file using a custom Perl script (Dietz et al. 2019) and mapped against a full coding sequence with BWA as described above. Diploid consensus sequences of the regions matching the reference were generated with samtools 1.6 (Li et al. 2009) and bcftools 1.6 (https://github.com/samtools/bcftools). The consensus sequences were further converted into alignments by another custom script (Dietz et al. 2019) to generate the individual gene alignments. This was followed by a gap removal from these alignments. In order to minimize the amount of missing data, which can potentially introduce biases, we generated two data sets: one with the loci that are present in at least 50% of the specimens (TE30) and another with loci that are present in all the specimens (TE60). These alignments were then manually checked in Geneious ver. 6.1.8 to make sure that there were no misalignments. Finally, individual loci alignments were concatenated for the initial phylogenetic analyses using FASconCAT-G (Kück and Longo 2014). Laboratory procedures and bioinformatics for ddRADseq followed Tahami et al. (2021).

### SNP calling

We extracted SNPs from the sorted BAM files generated during mapping in the TE30 dataset. SNPs were called using the samtools *mpileup* option piped together with the bcftools *call* option. Then filtering was done using bcftools, keeping one randomly chosen SNP per locus, filtering out indels and multiallelic SNPs. The resulting dataset contained a total of 3,164 SNPs, which were used for further analysis.

For ddRAD data, 18,383 SNPs were generated after the assembly and filtering where only those SNP sites for which data was present for at least 20 samples (*m*20) were taken into consideration.

### Phylogenetic analyses

For the target enrichment data, the best partitioning scheme for the concatenated dataset was found using PartitionFinder with parameter TESTMERGEONLY and rcluster algorithm (Lanfear et al. 2014), with the rcluster percentage set to 10, under the AICc criterion. The best partitioning scheme was then used as an input to set up a partitioned analysis in IQ-TREE ver. 2.0.3 (Chernomor et al. 2016; Minh et al. 2020b). We used the ultrafast bootstrap approximation with 1000 replicates (Hoang et al. 2018). To further reduce the risk of overestimating branch supports, the -bnni option was used. For ddRAD data, the TVM+F+I+G4 model of sequence evolution was used to reconstruct the ML tree in IQTREE using the ultrafast bootstrap approximation to compute 1000 replicates (Tahami et al. 2021). The resulting phylogenetic trees were visualised using FigTree (https://github.com/rambaut/figtree/releases) and rooted on *M. britomartis*. We compared the resulting trees using the *tipdiff* function in the R package treespace (Jombart et al. 2017). This function finds the number of differences in ancestry of the tips of two different trees with the same tip labels.

### Species tree analyses

Taking into consideration the gene tree-species tree discordance and incomplete lineage sorting that is common in datasets with multiple loci, we inferred a species tree using the summary-based coalescent method ASTRAL-III v. 5.7.4 (Zhang et al. 2018), which is statistically consistent under the multispecies coalescent framework. Model selection on each individual gene alignment from the TE60 dataset was performed using ModelFinder (Kalyaanamoorthy et al. 2017) and tree inference was done in IQTREE. The resulting output gene trees were used as an input for ASTRAL, which generated a species tree along with the quartet score. This score is the fraction of the induced quartet trees in the input set that occur in the species tree. The branch lengths of the ASTRAL species tree are in coalescent units and support values are given as local posterior probabilities (Zhang et al. 2018).

### Gene and Site concordance factor (gCF and sCF)

We calculated gene and site concordance factors (gCF and sCF) in IQTREE ver. 2.0.3 (Minh et al. 2020a) using the same set of gene trees used as an input for ASTRAL. gCF is calculated for each branch of a species tree as the fraction of decisive gene trees concordant with this branch and sCF as a fraction of decisive alignment sites supporting that branch (Minh et al. 2020a). The gene trees were compared against the species tree generated in IQTREE using a concatenation approach and various metrics were generated for the TE60 dataset. To better understand how concordance factors relate to each other and to the bootstrap values, resulting gCF and sCF values were plotted in R v. 4.0.3 (Lanfear, 2018).

### Population genetics

To get an initial idea of genetic clusters of related species present in the dataset, a principal component analysis (PCA) was carried out on the target enrichment SNP dataset and on all genotypes as well as unlinked SNPs obtained from ddRAD dataset using the *dudi.pca* function from the R package *adegenet* (Jombart and Ahmed 2011). We also performed a STRUCTURE analysis (Pritchard et al. 2000) to compare the admixture patterns inferred from ddRAD data and the target enrichment data. For this, our target enrichment SNP data in vcf format was converted to structure format (.str) using PGDSpider ver.2.1.1.5 (Lischer and Excoffier 2012). We tested 5 putative numbers of clusters, K=1-5, with 10 iterations for each K. To determine the optimal number of genetic clusters (K), we used the ΔK method in STRUCTURE HARVESTER (Evanno et al. 2005; Earl and vonHoldt 2012) with 500 000 generations for the Markov chain and a value of 100 000 as burn-in. The same parameters were used for the target enrichment and the ddRAD data set. We then aligned the cluster assignments of K=2, K=3 and K=4 across all the 10 replicates in CLUMPP (Jakobsson and Rosenberg 2007) and used DISTRUCT (Rosenberg 2004) to visualise the patterns of admixture from aligned clusters for both the datasets.

### Species delimitation

We performed a coalescent-based Bayes factor species delimitation implemented in SNAPP (Bryant et al. 2012) using the BFD* method (Leaché et al. 2014) on both target enrichment and ddRAD datasets. For ddRAD data, we used unlinked SNPs to test the species delimitation scenarios in SNAPP. We generated a subset of 30 taxa (Supplementary table S1) and tested 2 alternate scenarios. In the first scenario, the Balkan lineage of *M. athalia* was separated from the rest of the *M. athalia* and *M. celadussa* populations (Run C, table 3) and in the second scenario *M. athalia* (except for Balkan lineage of *M. athalia*) was lumped together with *M. celadussa* (Run D, table 3). The current taxonomy comprises 2 species – *M. athalia* (also including the Balkan lineage) and *M. celadussa*. For each scenario, a path-sampling analysis was carried out with 10 million generations. The run parameters including priors are described in the supplementary file. After the run for each of the alternate scenario and current taxonomy, SNAPP generated Marginal likelihood estimates (MLE). Bayes factor (Kass and Raftery 1995) is calculated as the difference between current taxonomy and each of the alternate scenarios, which is then used to assess the strength of species delimitation models. As per the BF scale, 0 < BF < 2 is not worth more than a bare mention, 2 < BF < 6 indicates positive evidence, 6 < BF < 10 represents a strong support, BF > 10 indicates decisive support. The calculation of Bayes Factors and ranking of different models followed the steps mentioned in Leaché and Bouckaet (2018).

**Table 2:**
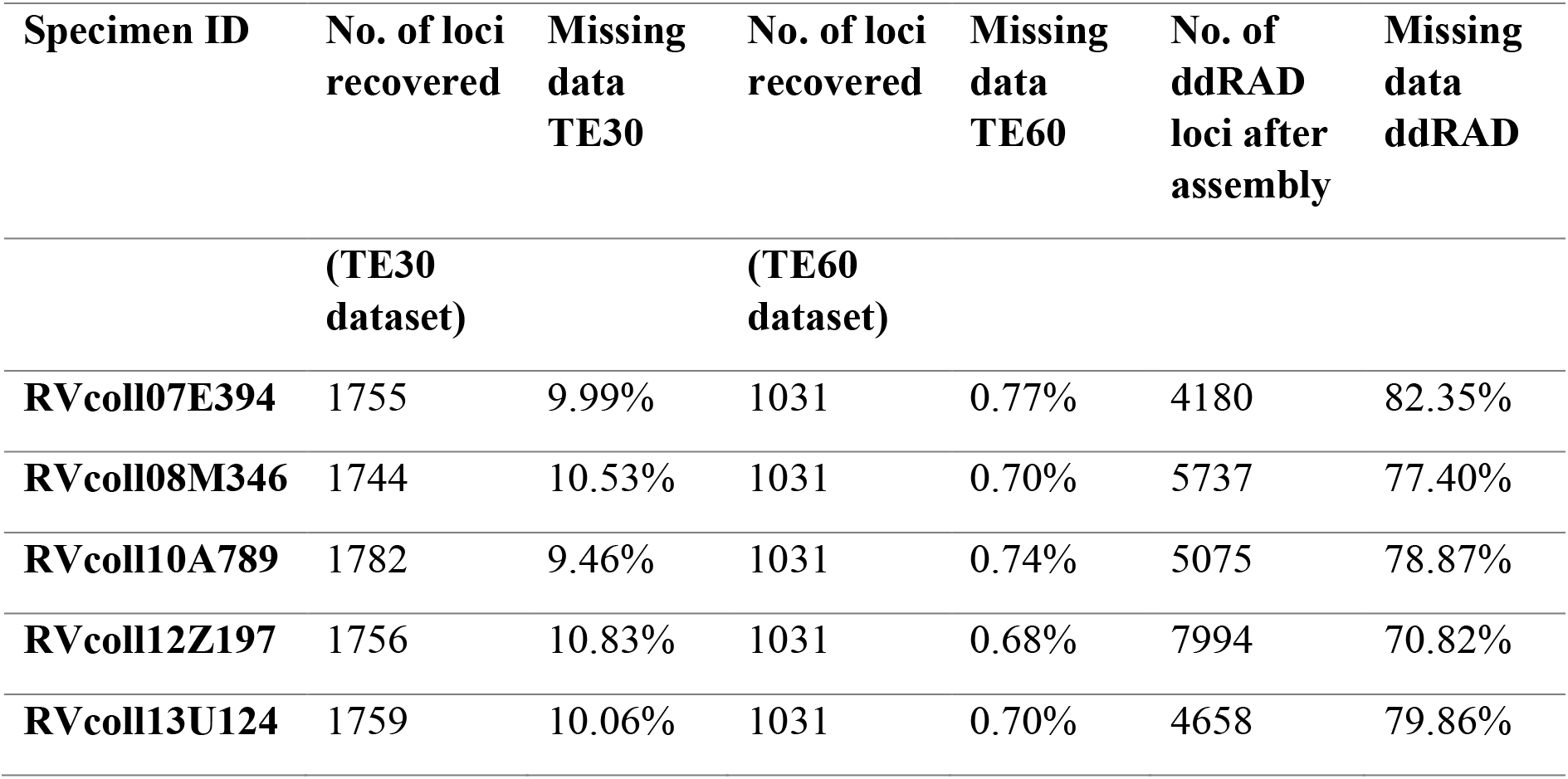
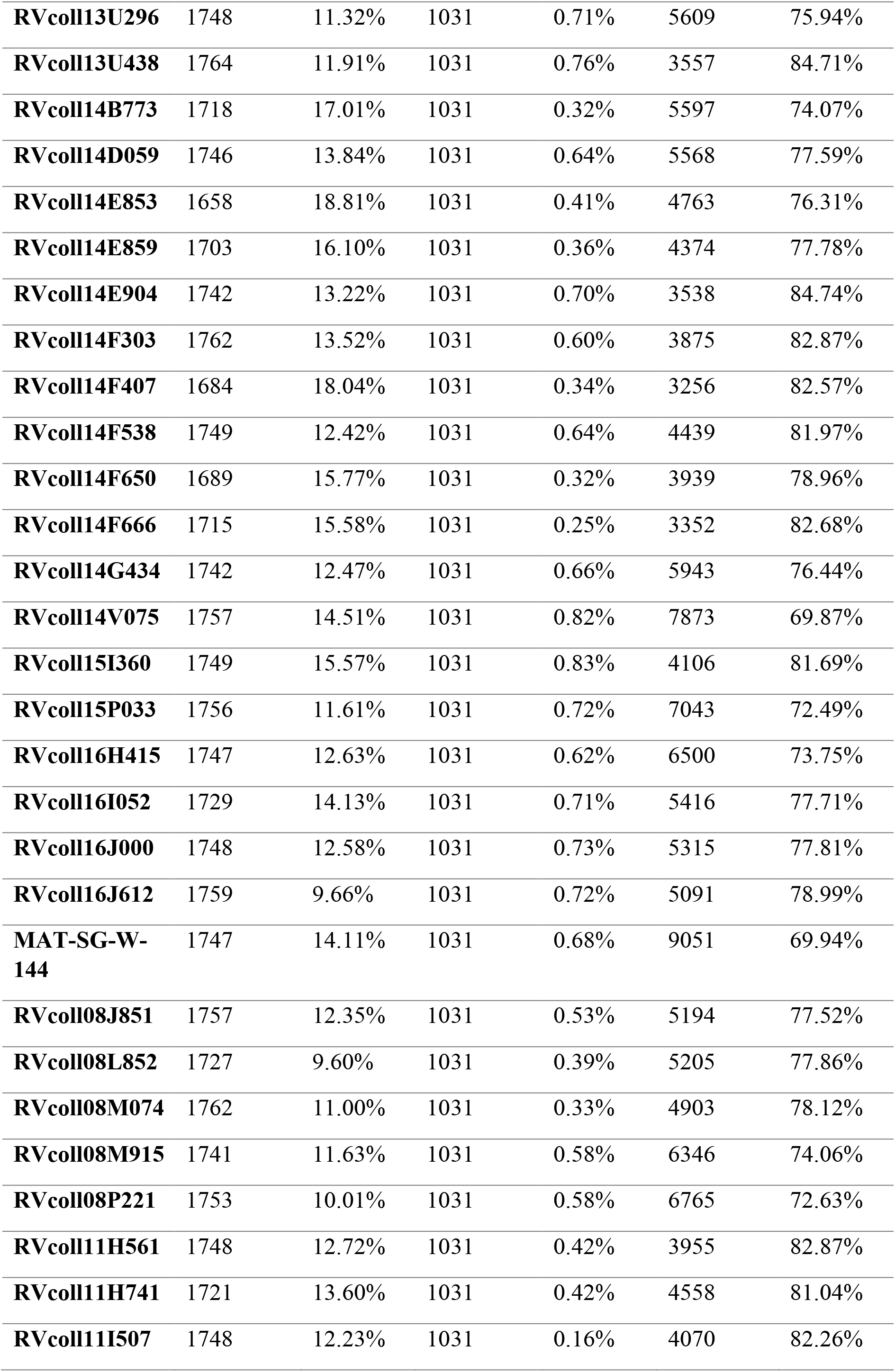
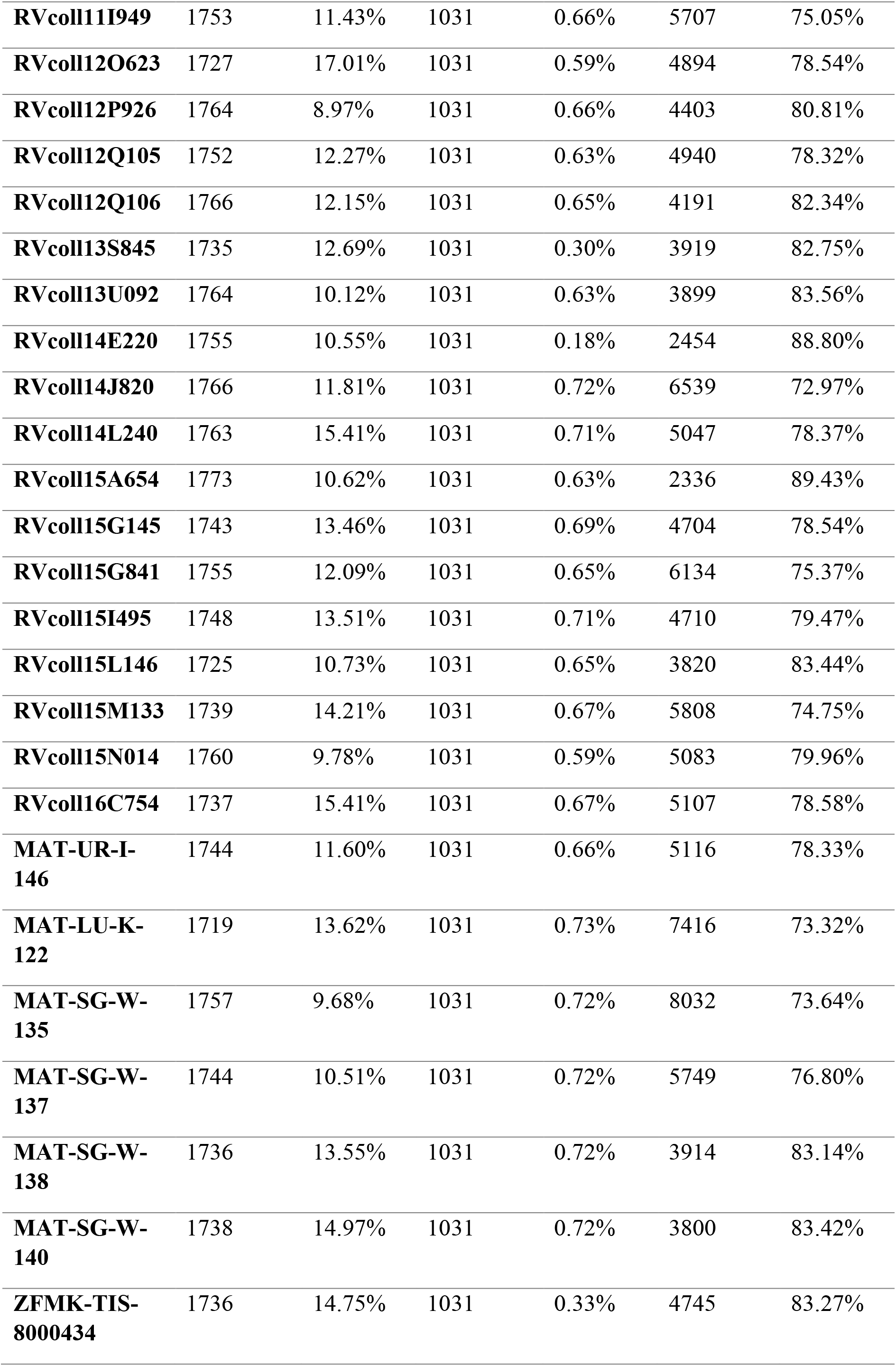
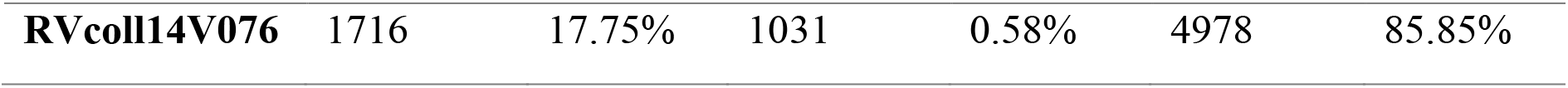
Overview of number of loci and missing data for the target enrichment and ddRAD datasets.

**Table 3:**
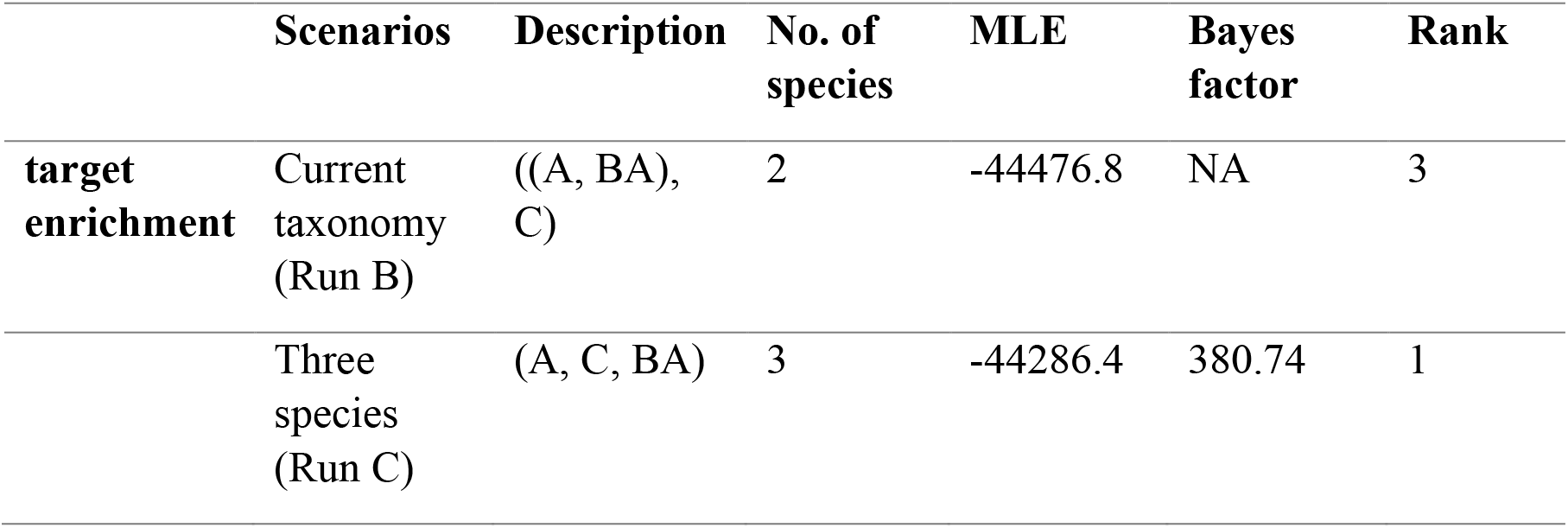
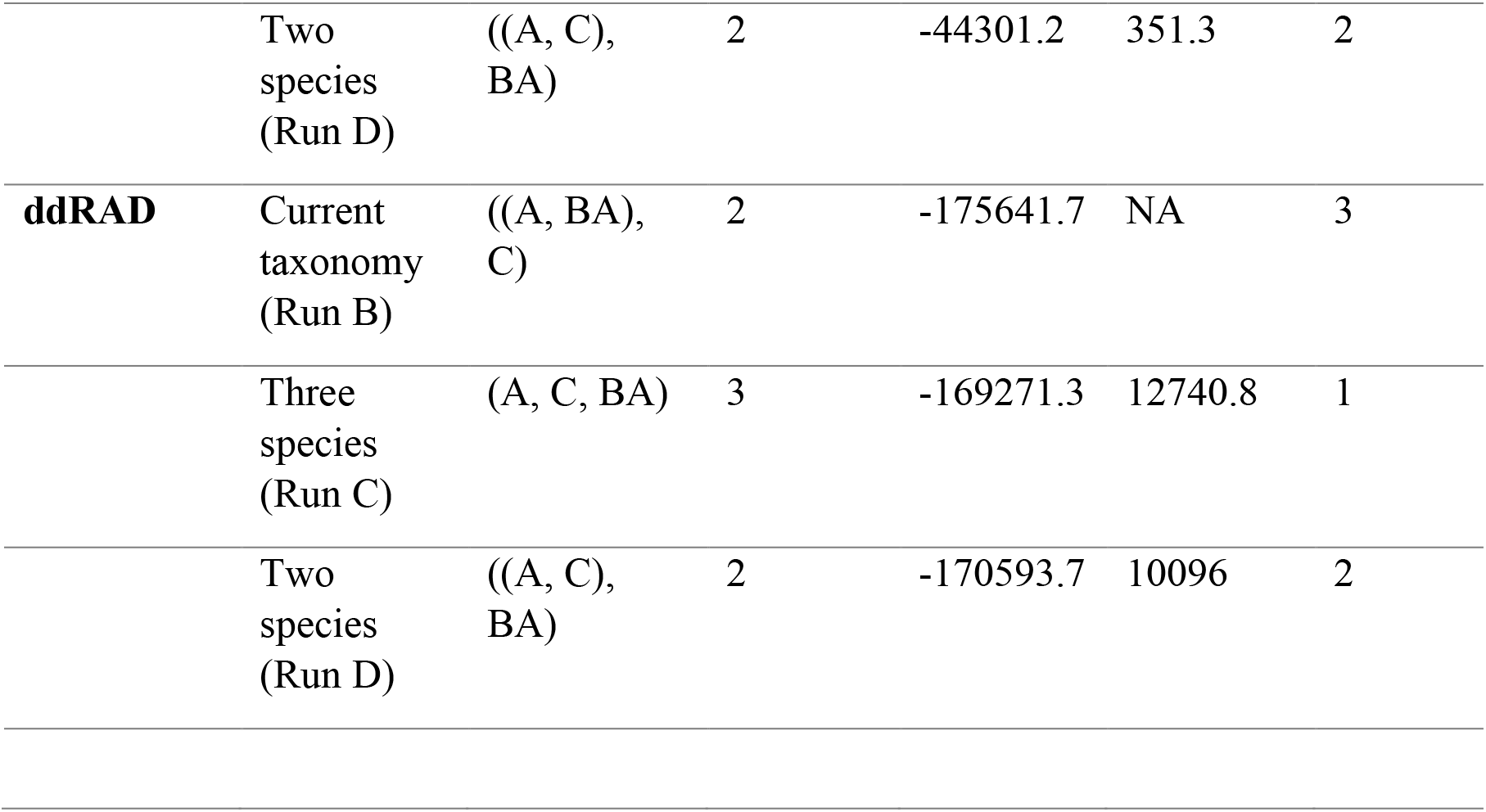
Comparison of different species delimitation models. Calculation of Bayes factor followed this tutorial - BFD-tutorial-1.pdf(netdna-cdn.com) A = *M. athalia* excluding Balkan lineage, BA = Balkan lineage of *M. athalia*, C = *M. celadussa* BF = 2*(MLE_1_ – MLE_0_), where MLE_1_ represents and alternate scenario and MLE_0_ represents current taxonomy.

|  | Scenarios | Description | No. of species | MLE | Bayes factor | Rank |
| --- | --- | --- | --- | --- | --- | --- |
| **target enrichment** | Current taxonomy (Run B) | ((A, BA), C) | 2 | -44476.8 | NA | 3 |
|  | Three species (Run C) | (A, C, BA) | 3 | -44286.4 | 380.74 | 1 |
|  | Two species (Run D) | ((A, C), BA) | 2 | -44301.2 | 351.3 | 2 |
| **ddRAD** | Current taxonomy (Run B) | ((A, BA), C) | 2 | -175641.7 | NA | 3 |
|  | Three species (Run C) | (A, C, BA) | 3 | -169271.3 | 12740.8 | 1 |
|  | Two species (Run D) | ((A, C), BA) | 2 | -170593.7 | 10096 | 2 |

In addition to the SNAPP analysis, we also tested the alternate species assignments using the *tr2* program, a multilocus species delimitation method that finds the best delimitation based on a distribution model of rooted triplets (Fujisawa et al. 2016). We used rooted individual gene trees from the TE60 dataset as input. The two models we tested, which correspond to the two-species and three-species hypotheses respectively, were compared against a null model (which assumes the presence of a single species) based on - log(likelihood) scores.

## RESULTS

### Overview of the datasets

After the read filtering step, an average of 1.7 million reads were recovered across all the specimens for the target enrichment dataset (Table 1). For the TE30 dataset, the average number of informative loci retained was 1,743 (SD= 21.9, Table 2), with an average amount of missing data of 12.79% (SD=0.023, Table 2). From this dataset, we further removed the loci with zero or very few variable sites (number of variable sites = 0, 1 or 2). The final average number of loci retained was 1,582. For the TE60 dataset, 1,031 loci were retained with the average amount of missing data dropping to 0.60% (Table 2). After removing the loci with zero or few variable sites, 1,002 loci were retained. For the ddRAD dataset, the average number of loci retained after assembly was 5,071 (Table 2), but the percentage of missing data was much higher than for the target enrichment dataset (avg. 78.9%, Table 2).

### Target Enrichment Phylogenetics

For the sake of communication purposes, we labelled our specimens according to information provided by their mitochondrial COI barcodes. Two separate trees were generated for the two target enrichment datasets, TE30 and TE60. In both cases, six specimens of *M. athalia* originating from the Balkans (atha14F407, atha14F660, atha14F666, atha14B773, atha14E859, atha14E853) were recovered as a distinct lineage (Balkan clade) sister to the rest with 100% bootstrap support (Fig. 1 and Fig. 3a). However, this Balkan clade did not include all the specimens from Balkan region that we analysed. The remaining specimens of *M. athalia* were found to be paraphyletic with respect to *M. celadussa* (BS 100, Fig. 1). Within this group, another set of six *M*. *athalia* individuals from Balkans (atha14V075, atha08M346, atha14F303, atha14F538, atha14D059, atha14G434) formed a clade sister to the rest (BS 99 in TE60, BS 100 in TE30). We did not find a clear separation of specimens from the contact zone or of specimens with intermediate genital characters (from morphometric analyses done by Tahami et al. (2021)).

**Figure 1:**
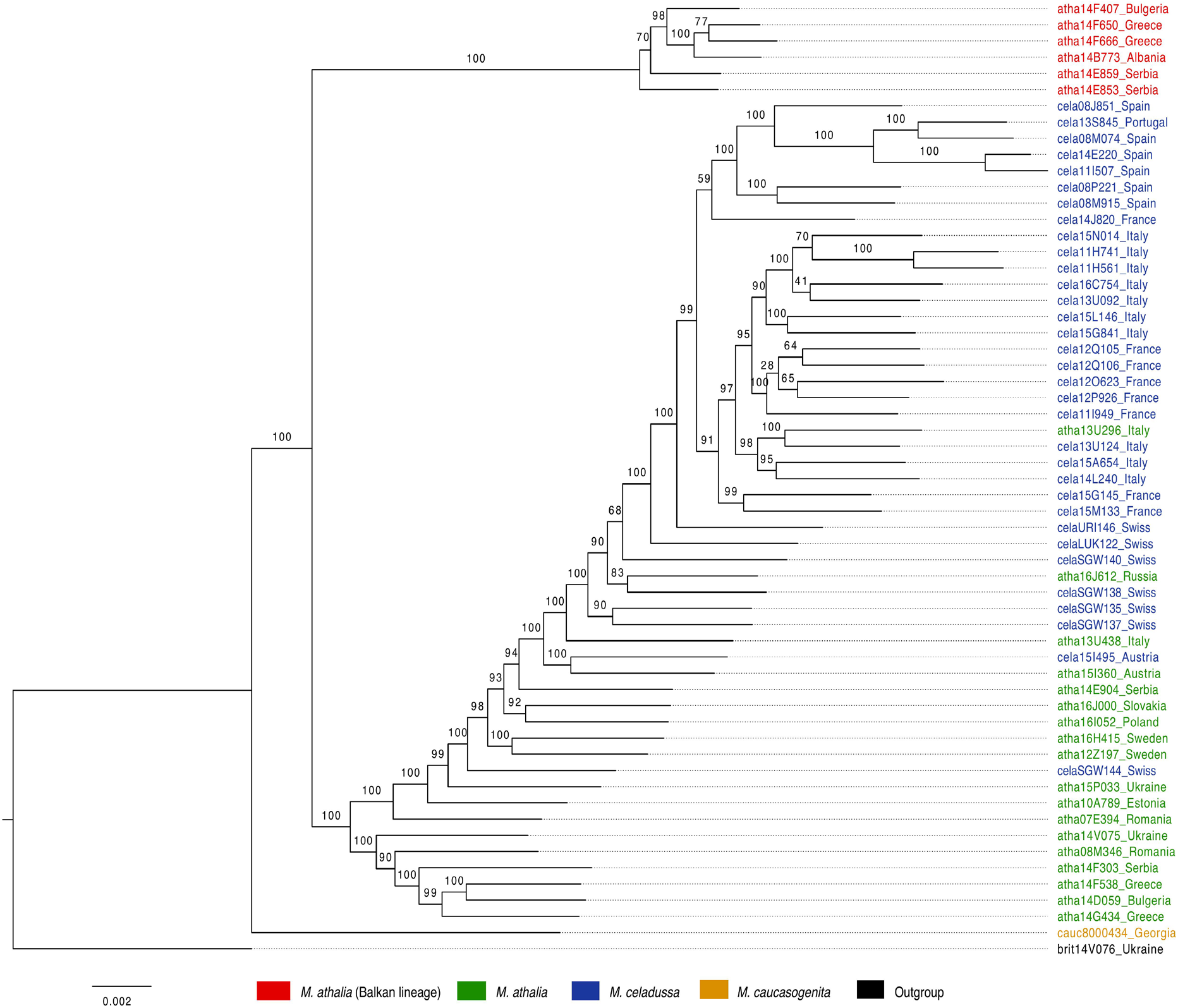
Maximum Likelihood (ML) phylogenetic tree based on the target enrichment TE30 dataset rooted on *M. britomartis*. Numbers on the branches indicate bootstrap values calculated based on 1000 replicates.

**Figure 2:**
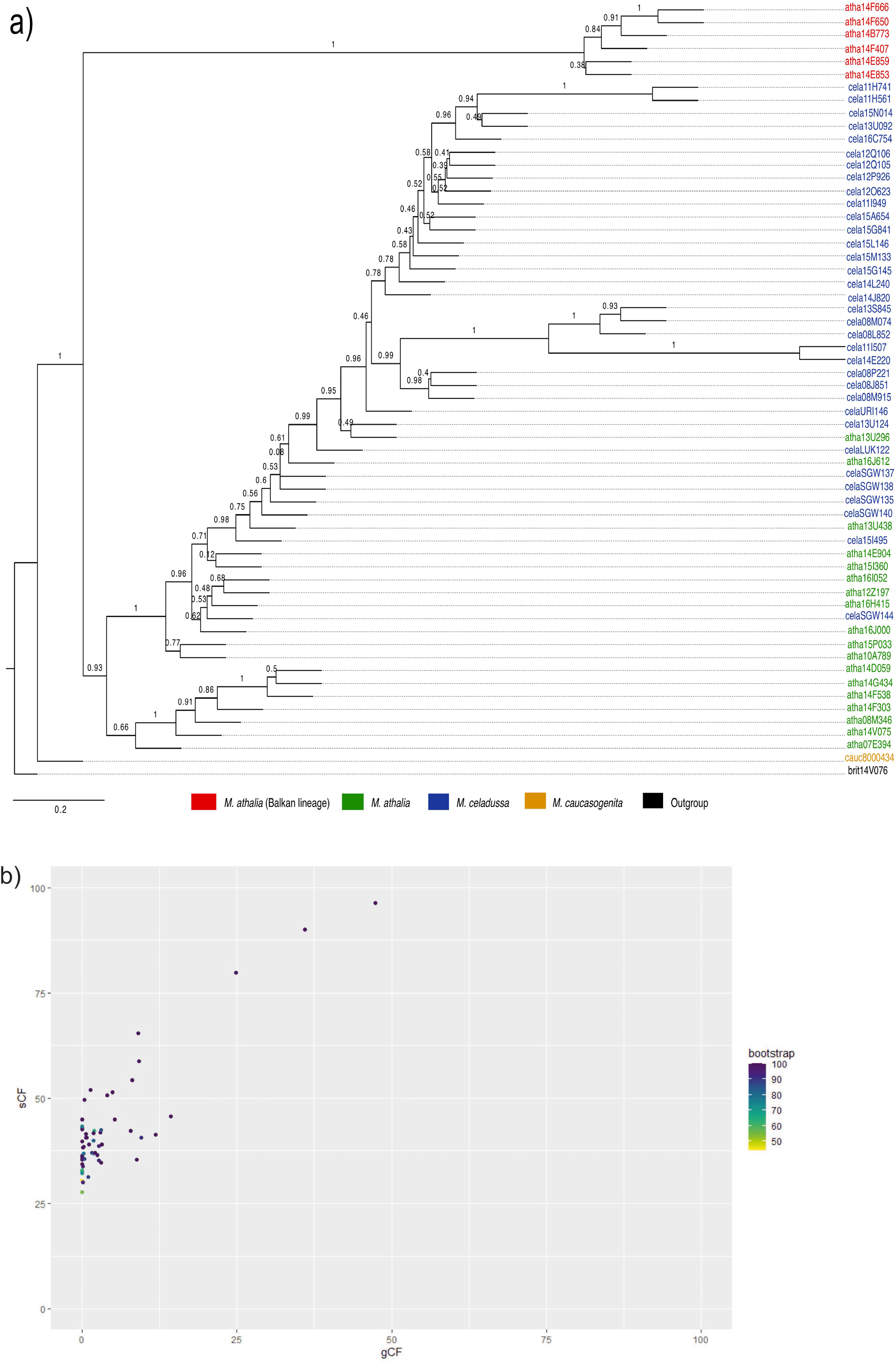
(a) ASTRAL species tree based on the target enrichment TE60 dataset and rooted with *M. britomartis*. Numbers on the branches are quartet support scores. (b) sCF vs. gCF for the TE60 dataset.

**Figure 3:**
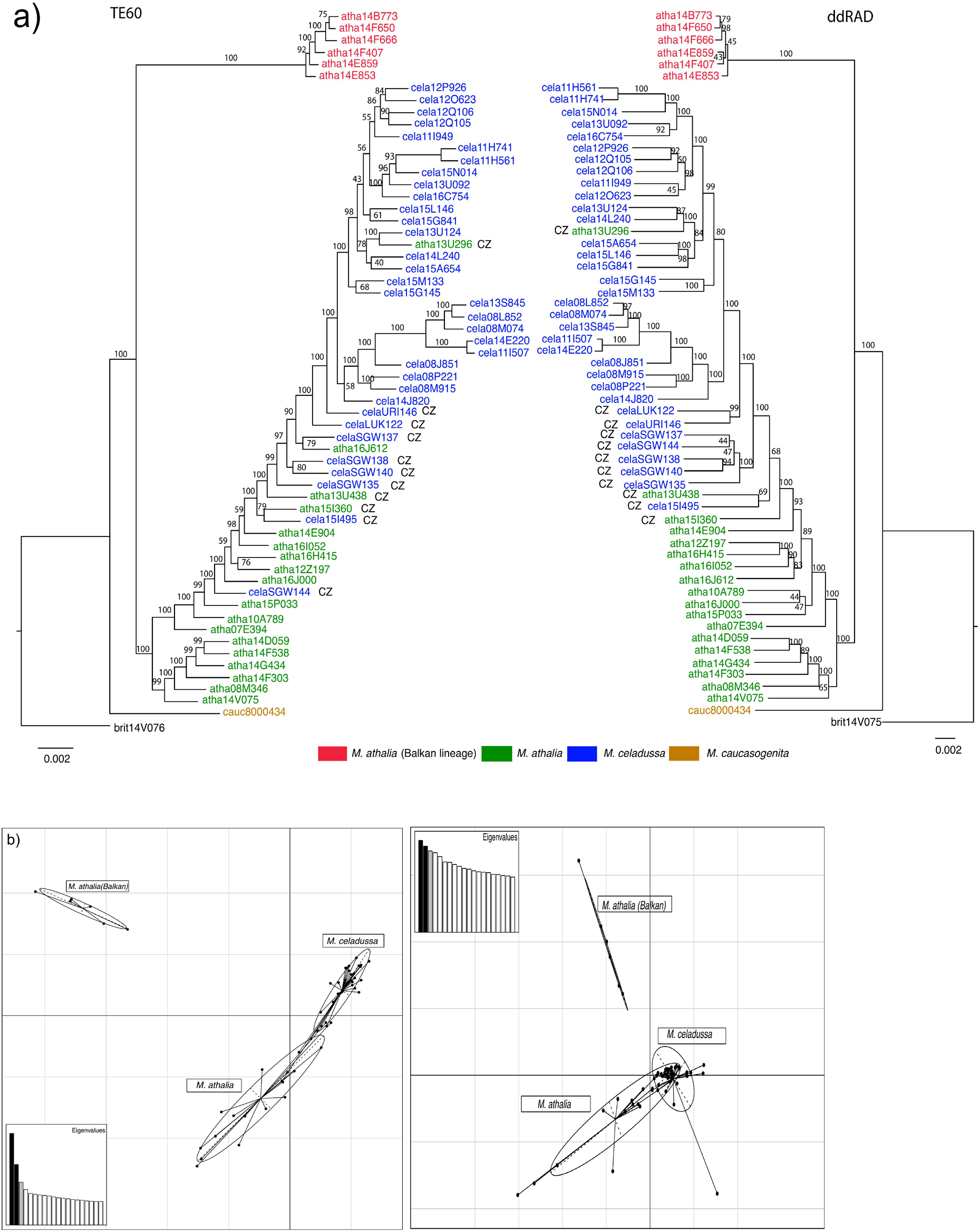
(a) Comparison of maximum-likelihood phylogenetic trees obtained from the TE60 dataset (on the left) and ddRAD dataset (on the right). Both trees are rooted with *Melitaea britomatris.* Numbers on the branches indicate bootstrap values calculated based on 1000 replicates. Contact zone specimens are indicated with abbreviation ‘CZ’ next to them (b) PCA plots for target enrichment SNP dataset (on left, PC1 and PC2 with variances 6.93% and 4.57% respectively) and ddRAD dataset (on right, PC1 and PC2 with variances 3.46% and 3.23% respectively).

### ASTRAL species tree and concordance factors

The general patterns found in the ASTRAL species tree were congruent with the maximum likelihood (ML) phylogenetic analyses in IQ-TREE, with the Balkan *M*. *athalia* clade being supported with local posterior probability of 1, and a clade containing paraphyletic *M*. *athalia-M. celadussa* grade having local posterior probability of 0.93 (Fig. 2a). The same set of six *M. athalia* individuals as in the ML analyses grouped together at one end of this clade (posterior probability 0.66). The scoring of this tree gave a final quartet score of 235095870, and only 48.11% of all the quartet trees could be found in the species tree (normalized quartet score of 0.48) indicating high gene tree discordance.

In the species tree obtained from the IQ-TREE concatenation approach (supplementary figure S1), higher values for both concordance factors were mainly observed for the Balkan *M*. *athalia* clade. For the paraphyletic *M*. *athalia-M. celadussa* grade, and branches within it, gCF values were lower even though sCF values were (moderately) on the higher side. The same was observed when plotting sCF vs. gCF (Fig 2b), where most of the branches fall on the lower side for gCF values (near 0 and between 0 to 25) and moderately high side for sCF values (between 25 to 75), while a few branches had higher values of both concordance factors (gCF above 25 and sCF above 75).

### Comparison – Phylogenetics

We compared a phylogenetic tree obtained from the TE60 dataset with a phylogenetic tree generated using ddRAD data, excluding the taxa that are not present in the target enrichment dataset (Fig. 3a). The general phylogenetic relationships were the same in both datasets. Both methods suggested the presence of a distinct Balkan lineage, which in both cases included the same six samples (Fig. 3a, individuals indicated in red color). The near-monophyly of *M. celadussa* is confirmed in both trees, with the exception of specimens from the contact zone. A clade containing another set of six *M*. *athalia* individuals from the Balkans, which is sister to the paraphyletic *M*. *athalia*-*M*. *celadussa* grade had a rather low bootstrap support (BS=65) in the ddRAD analyses, and higher support in the target enrichment analyses (BS=99). In both cases, the individuals identified by biodecrypt as belonging to the contact zone formed part of the paraphyletic *M*. *athalia*-*M*. *celadussa* grade. From the comparison using *tipdiff* (supplementary figure S2), it can be seen that the number of differences between the two trees for the six diverging Balkan samples is zero, whereas there are two to four differences in the *M*. *athalia* clade that form sister to the paraphyletic grade, and 42-43 differences for the rest of the taxa from the paraphyletic *M*. *athalia*-*M*. *celadussa* grade.

### Comparison – Population genetics

The PCA plots for both target enrichment and ddRAD showed a clade of *M. athalia* from the Balkan, that was found to be genetically distinct from all other studied specimens, including some other specimens from Balkan (Fig. 3b). The clusters for the rest of the *M. athalia* and *M. celadussa* were found to overlap to some extent in PCA plots for both datasets (Fig. 3b). The analysis of genomic admixture using STRUCTURE produced the highest likelihood estimate for K=2 clusters for the ddRAD dataset, consistent with the results obtained by Tahami et al. (2021) and K=3 for the target enrichment dataset (Supplementary figure S4). To better visualise the patterns of admixture spatially, we also mapped the membership coefficient matrix obtained from Bayesian clustering at K=3 for target enrichment to the geographic coordinates plotted on a map in R v 4.0.4 (R core team, 2021). The comparison of STRUCTURE barplots from K=2 to K=4 for both datasets (Fig. 4a) shows the distinctiveness of the Balkan *M. athalia* clade. These contrast with the presence of admixture in many of the *M. athalia* specimens (both in the contact zone and notably far from it) and in several *M. celadussa* individuals from the contact zone. This pattern is also evident from the geographic distribution of genomic admixture (Fig. 4b), where the Balkan clade, some *M. athalia* (in the Balkans, Eastern Europe and N. Scandinavia) and the *M*. *celadussa* from south-western Europe (Spain, France and most of Italy) are shown to have pure gene pools (green, orange and pink coloured pies on the map, respectively), whereas the individuals from the contact zone and from a wide area in central Europe showed considerable genomic admixture (Fig. 4b).

**Figure 4:**
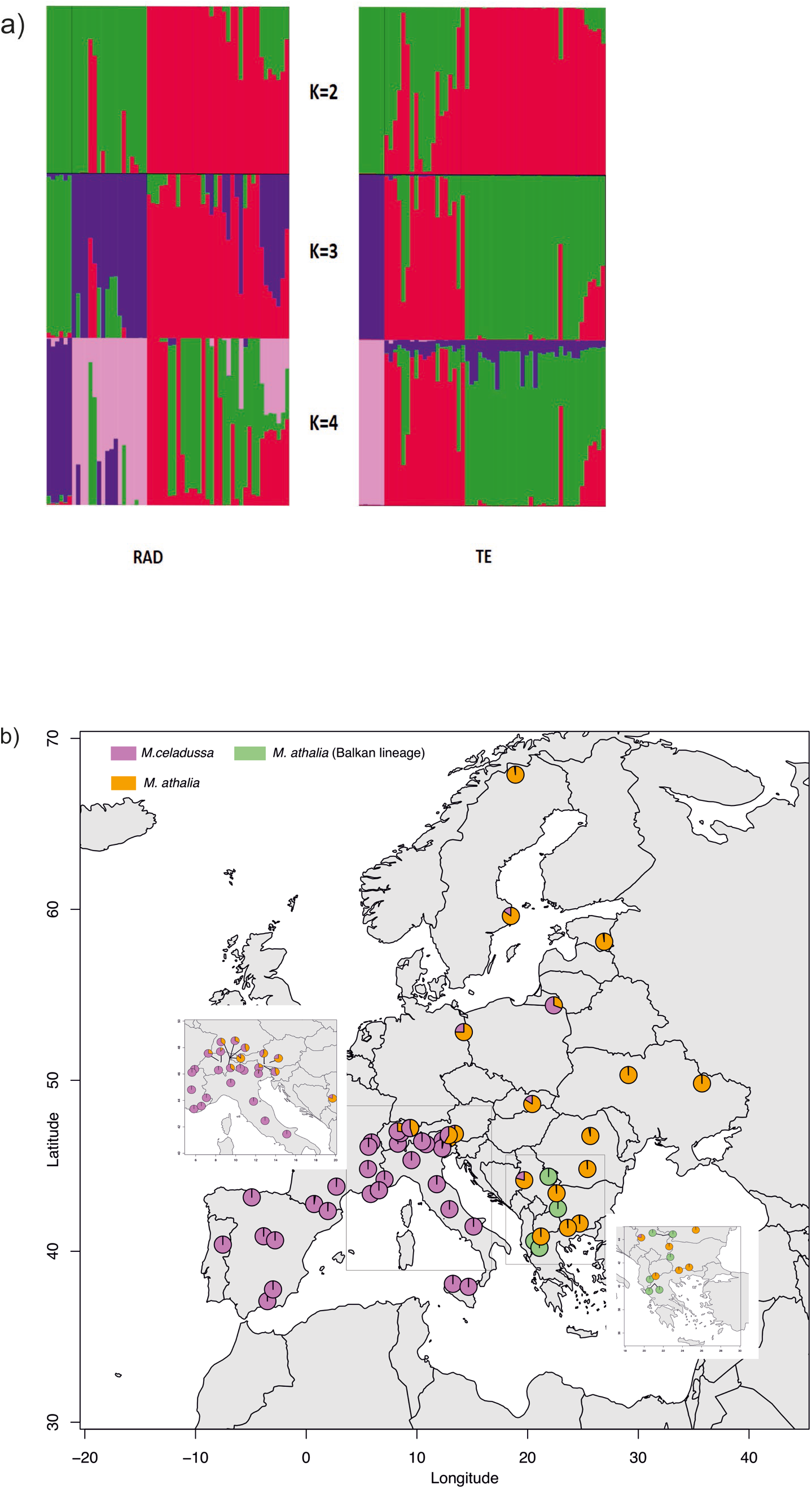
(a) STRUCTURE analysis of *M. athalia*-*M. celadussa* for ΔK=5 for the target enrichment dataset (right) and ddRAD dataset (left). The aligned barplots show cluster assignments at 2-4, from top to bottom. (b) Pie charts based on membership coefficient matrix (q-matrix) at K=3 for the target enrichment dataset mapped on geographic coordinates.

### Comparison - Species delimitation

Based on Marginal Likelihood Estimates (MLE, Table 3), the three-species scenario achieved the highest rank among the two scenarios tested for both datasets (Table 3). However, as per BF scale, the BF values for both two-species (Run D, lumping *M*. *athalia* except Balkan lineage and *M*. *celadussa* together) and three-species scenarios (Run C, Balkan *M*. *athalia, M. athalia* and *M*. *celadussa*) were greater than 10, hence were concluded as decisive for both target enrichment and ddRAD datasets (Table 3). From the tr2 analysis, negative log likelihood values −3659835.67, −2101731.61 and −123421.81 were calculated for the null model, model1 (with two species) and model2 (with three species) respectively. Thus, the likelihood is distinctly higher for the three species model.

## DISCUSSION

Using a parapatric pair of *Melitaea* butterflies as a model system, we compared two genomic approaches for their utility in elucidating patterns of genetic structure and admixture, and investigated whether genome-wide datasets could provide means for well-informed and robust species delimitation under parapatry. Both sequence capture and RADseq have been independently explored for species delimitation before (Smith et al. 2014; Pante et al. 2015; Gueuning et al. 2020), but to the best of our knowledge, our study is among the first to focus on species delimitation in a parapatric system (Linck et al. 2019 being a rare exception). Additionally, only few studies have compared two genomic approaches in the same study system (Leaché et al. 2015; Harvey et al. 2016; Manthey et al. 2016), all of them utilising UCEs as a primary sequence capture method. Consequently, there has been little discussion over the benefits and drawbacks of various genomics methods in species delimitation.

In our study system, both the maximum likelihood inference and ASTRAL species tree inference recovered the same set of major clades in both the target enrichment and ddRAD approaches, thus providing a consistent phylogenetic picture for the group. *M. celadussa* was long considered as a subspecies of *M. athalia* (Higgins 1955) but was recently given a full species status (Leneveu et al. 2009; Wiemers et al. 2018). We recovered *M. celadussa* sensu Leneveu et al. (2009) as a near-monophyletic group with few exceptions and most of *M*. *athalia* as a paraphyletic grade with respect to *M. celadussa*. Specimens for which intermediate genital characters were reported (and which according to DNA barcodes are *M. celadussa*) were also part of this paraphyletic *M. athalia*-*M. celadussa* grade. A higher discordance among gene trees was observed for the *M*. *athalia*-*M. celadussa* grade as per gCF analyses, the conflict possibly arising due to introgression of *M. celadussa* genes into *M. athalia*. Also, most differences between ddRAD and target enrichment trees were found for this grade as per *tipdiff* plot. Comparison of patterns of admixture using STRUCTURE analyses revealed different optimum K values for the two methods. However, the PCA plots looked largely similar, with a distinct Balkan lineage cluster and two overlapping clusters of *M. athalia* and *M. celadussa*. The barplot for STRUCTURE analyses at K=2 for ddRAD and K=3 for target enrichment, as well as the geographic distribution of genomic admixture (for target enrichment K=3), show patterns consistent with those observed by Tahami et al. (2021), with Balkan *M*. *athalia* and south-western European *M. celadussa* having non-admixed gene pools and a high admixture in individuals from the contact zone and in *M. athalia* in C. Europe. One notable difference is that, in the case of the target enrichment dataset, the clusters tended to be purer. For example, the Balkan clade and *M. celadussa* specimens from south-western Europe are clearly shown as non-admixed in the target enrichment-based barplot and admixture pie plots mapped on geographic coordinates. This could be attributed to the low amount of missing data, and therefore less noise in the target enrichment dataset (Table 2). For the species delimitation analyses, both the three-species and two-species scenarios received a decisive support by target enrichment and ddRAD, suggesting that both datasets supported each of the two scenarios equally. However, testing of alternate species assignments using tr2 delimitation for target enrichment TE60 data favoured the 3-species hypothesis over the 2-species one.

From our results, the three-species scenario seems to be more strongly supported than the two-species one. Based on ddRAD data, Tahami et al. (2021) hypothesized that the taxa examined here could have originated from three main refugia in Europe along several glacial cycles: Iberia and Italian Peninsula for *M. celadussa*; the Balkan Peninsula for the Balkan lineage; and an eastern refugium for non-admixed *M. athalia*. They also hypothesized that the observed patterns could have resulted from adaptive introgression of a set of *M. celadussa* genes eastward into *M. athalia*. Based on the results from current study, we see that two main genetically distinct entities are identified (Balkan lineage of *M. athalia* and rest of the *M. athalia* and *M. celadussa* specimens) which agrees with the two-species scenario. But *M. athalia* and *M. celadussa* are mainly identified as separate species by large distribution, genital differentiation and mitochondrial and nuclear genetic separation. In addition to that, there is an unexpected split between two lineages of *M*. *athalia* where the genetically diverging Balkan lineage stays distinct from rest of the *M. athalia* from this region despite geographical proximity, which could be possible explanation the three-species scenario.

### Target enrichment vs. ddRAD – Benefits and pitfalls

In target enrichment methods the sequencing efforts are concentrated on a pre-defined set of loci, and therefore high coverage and little missing data is expected (Mayer et al. 2021). Different target enrichment methods mainly differ in the gene regions they target and, as researchers can select loci at desired levels of divergence (Banker et al. 2020), they have been shown to be generally efficient in elucidating affinities at both deep and shallow phylogenetic scales (Hamilton et al. 2016; Espeland et al. 2018; Bagley et al. 2020).

Using a recently developed target enrichment kit that targets about 2900 CDS regions (Mayer et al. 2021), we were able to recover many gene loci across the individuals even after stringent filtering criteria (1002 gene loci for TE60) which shows the capture experiment to be successful. We found that the number of loci captured was sufficient to uncover the phylogenetic relationships of *Melitaea*-irrespective of whether loci were concatenated and assumed to have the same genealogical history, or if they were allowed to have independent histories and analysed within a summary coalescent framework using ASTRAL. This demonstrates that the target enrichment loci contain sufficient phylogenetic information that could be useful to delimit closely related taxa. Also, the amount of missing data in both target enrichment datasets was very low compared to the ddRAD dataset.

RADseq methods are inexpensive and quick compared to target enrichment and do not need genomic resources to design probes. However, they are most useful at shallow scales of divergence, as the number of homologous loci obtained decreases with phylogenetic distance (Wagner et al. 2013; Lee et al. 2018). Furthermore, the locus dropout effect also increases with divergence, leading to non-random patterns of missing data (Arnold et al. 2013). RADseq methods also require high quality material, i.e., better preserved tissue to successfully extract enough amounts of DNA, while it is possible to recover genomic data from old museum material using target enrichment (Call et al. 2021; Mayer et al. 2021). In addition, probes used for hybrid enrichment can enrich regions containing up to 30% mismatches to the probe sequences and thus can efficiently capture the loci also if the variation is higher, while ddRAD requires exact matches of short k-mers for restriction enzymes to make cuts consistently across individuals (Banker et al. 2020).

It has been suggested that the method of data collection should not influence the results of phylogenetic analyses at intermediate levels of divergence (Manthey et al. 2016). From our comparison of target enrichment and ddRAD, this should also be the case at an alpha taxonomic level. Therefore, the choice of method to tackle species delimitation at such levels could be entirely up to the researcher based on resources available and quality of the samples. Here, some practical aspects are worth taking into consideration: target enrichment is generally more expensive than RADseq because of the costs associated with the library preparation and purchasing enrichment probes (Harvey et al. 2016). The target enrichment laboratory work tends to be slower due to additional hybridization and enrichment steps although with the newer kits it is now significantly faster. However, as the loci obtained from RADseq are random and not known, different RAD datasets are hard to combine and are not comparable due to their unique nature. Different RAD datasets are also likely to be composed of loci with different levels of conservativeness (Lee et al. 2018). Target enrichment has an important advantage in terms of cross-compatibility, since the targeted loci remain fixed. As the loci being recovered are already known in target enrichment and usually have low levels of missing data, this could be an advantage in identifying a universal set of markers useful for species delimitation. It would also enable a comparison across different species groups, as well as combining different datasets, and hence provide means for standardized delimitation of e.g. allopatric populations and other settings where delimitation of species is inherently difficult.

Recently, the use of a standard set of nuclear markers for species delimitation and identification was proposed (Eberle et al. 2020), which can be studied in and compared across all animals and would allow disentangling recently diverged lineages. This idea was tested by Dietz et al. (2021) using empirical data from several metazoan lineages. They found that so-called ‘Universal Single-Copy Orthologs’ (USCOs) performed better than DNA barcodes to delimit closely related species, irrespective of the assembly approach or tree reconstruction method used (Dietz et al. 2021). A similar idea of having a unified set of loci for phylogenomic and population genetic studies has also been explored by Singhal et al. (2017), but exclusively for squamates. Basing species delimitation on a standard set of molecular markers would be an important step towards a more stabilized taxonomy, particularly under conditions where delimitation is bound to be arbitrary. This concerns the delimitation of allopatric and parapatric populations, as well as asexual strains, but would likely turn out to provide efficient means for a delimitation in all settings.

### The conundrum of parapatric species delimitation

Finding universally applicable criteria for species delimitation is rendered extremely challenging particularly due to the complexity of biological systems (Eberle et al. 2020) and the semantics around the meaning of the term ‘species’ (de Queiroz 2007). Different species concepts emphasize different properties of biological diversity, such as temporal and spatial aspects of variability, capacity for reproduction, type of reproduction, reciprocal monophyly, diagnosability and ecological uniqueness. The complexity of biological diversity is a central cause for the amplitude of species definitions that have been proposed. Attempts of considering many or all of these aspects inevitably lead to broad and hence vague definitions of species (e.g. the General Lineage Concept (de Queiroz 1999)). We find reaching a consensus over the definition of species not foreseeable and hence do not discuss this further here, but we are slightly more optimistic that the disagreement over the epistemological aspects of species delimitation could be overcome by adopting efficient and standardizable approaches such as those presented by Eberle et al. (2020) and Dietz et al. (2021).

Parapatric taxa constitute an unusually complex case for species delimitation, because they typically show more or less limited, but frequent, hybridization in the contact zone (Hewitt 1988; Bull 1991) and genetic admixture that may extend far beyond the zone of contact (Osada et al. 2010; Johnson et al. 2015). Parapatric taxa tend to be morphologically and ecologically very similar, and morphologically intermediate individuals may occur (Saino and Villa 1992; Guiller et al. 2017; Slender et al. 2017). From a temporal perspective, parapatric systems are presumably usually young – although initial differentiation may be much older (Ebdon et al. 2021) – and may have undergone either slow merging of populations or increased differentiation, the latter being promoted by disruptive selection and reinforcement (Barton and Hewitt 1985). These complexities also characterize our study system, which we assume is resulting from a postglacial secondary contact of populations differentiated in two refugia during the last glaciation, and possibly in earlier ones (Tahami et al. 2021).

As the delineation of parapatric taxa is inherently arbitrary and largely subjective, delimiting admixing parapatric taxa in a consistent manner would promote taxonomic stability. Optimally, delimitation should be based on broad genomic data, because it is little affected by choice of marker and provides sufficient data to obtain a proper understanding of evolutionary relationships and admixture between the populations. Generating a species list at global, continental or local scale for any group of organisms would require a widely accepted pragmatic solution for identifying parapatric taxa either as infraspecific or as full species. Parapatric systems are also characterized by notable discordance between different types of characters across individuals. Such discordance, both mitonuclear and mito-morphological, was also observed in *Melitaea*. Keeping in mind that admixing taxa or hybrids cannot be assigned to species by definition and that around the hybrid zone most specimens might fall into this category, we would need to identify which parameters are most relevant to the formation of evolutionary distinct units (frequency of hybrids, mitonuclear discordance, presence and frequency of morphological intermediates, distance from the contact zone where introgressed specimens can be found). A drawback of considering parapatric species as infraspecific is that this might prevent their recognition as biological entities worth of protection in countries where conservation legislation does not recognize the value of taxa below species level. However, the same holds true at all levels of biodiversity, including an uncountable number of genetically unique populations whose recognition as valuable units of biodiversity is seldom legally acknowledged.

## CONCLUSIONS

The emergence of high-throughput genetic tools is revolutionizing systematic research at all phylogenetic levels. Many platforms and protocols to generate genomic-scale datasets are available, each with their specific strengths and shortcomings. This poses a question: which particular method provides the best means to address the research question at a given phylogenetic level? In this work, we focused on answering this question at the interface of population and phylogenetic levels, using a particularly challenging system of two parapatric butterfly species with frequent introgression and widespread admixture as a model. While both ddRAD sequencing and target enrichment methods provided largely congruent pictures of the evolutionary history of our study system, the latter has the benefit of providing higher levels of scalability. Compared to RAD datasets, sequence capture is characterized by a dramatically lower degree of missing data due to a smaller locus dropout effect and sequencing of a pre-defined set of loci at a high coverage. Presently, there is no consensus over the principles for delimitation of parapatric taxa. We argue that this is inherently based on arbitrary criteria and reaching a widely accepted consensus is desperately needed. As patterns of admixture like those observed here are more a rule than an exception in parapatric systems, we propose regarding admixing parapatric taxa in a consistent way. Considering them regularly either as subspecies or full species would promote taxonomic stability, but both solutions would also be characterized by some shortcomings.

## DATA ACCESSIBILITY

All raw sequence data are archived on the NCBI Sequence Read Archive under BioProject ID PRJNA802033.

## SUPPLEMENTARY MATERIAL

Data to be deposited on dryad

## FUNDING

This work was supported by academy of Finland (grant no. 314702) to MM. RV was supported by Grant PID2019-107078GB-I00 funded by MCIN/AEI/ 10.13039/501100011033. VD was supported by the Academy of Finland (Academy Research Fellow, decision no. 328895).

## ACKNOWLEDGEMENTS

The authors would like to thank Sandra Kukowka for her help in the lab work. We would like to acknowledge CSC–IT Center for Science, Finland, for providing computational resources. We would also like to thank Tomochika Fujisawa for helpful discussion on tr2 analysis.

## Notes

### Competing Interest Statement

The authors have declared no competing interest.

