## Supplementary file for "Delimiting Continuity: Comparison of Target Enrichment and ddRAD for Delineating Admixing Parapatric *Melitaea* Butterflies"

**Table S1**: A list of specimens included in the SNAPP analysis

| **Specimen ID** | **Clade** |
| --- | --- |
| atha07E394 | athalia (A) |
| atha14E904 | athalia (A) |
| atha14B773 | Balkan athalia (BA) |
| atha13U296 | athalia (A) |
| atha10A789 | athalia (A) |
| atha14G434 | athalia (A) |
| atha12Z197 | athalia (A) |
| atha16J612 | athalia (A) |
| atha14E859 | Balkan athalia (BA) |
| atha14F407 | Balkan athalia (BA) |
| atha15I360 | athalia (A) |
| atha14E853 | Balkan athalia (BA) |
| atha14V075 | athalia (A) |
| atha16J000 | athalia (A) |
| atha14F650 | Balkan athalia (BA) |
| cela13U124 | celadussa ( C ) |
| cela15G841 | celadussa ( C ) |
| celaLUK122 | celadussa ( C ) |
| cela13S845 | celadussa ( C ) |
| cela11H561 | celadussa ( C ) |
| cela15G145 | celadussa ( C ) |
| celaSGW135 | celadussa ( C ) |
| celaSGW138 | celadussa ( C ) |
| cela11I949 | celadussa ( C ) |
| cela08J851 | celadussa ( C ) |
| celaURI146 | celadussa ( C ) |
| cela15I495 | celadussa ( C ) |
| cela15L146 | celadussa ( C ) |
| celaSGW144 | celadussa ( C ) |
| celaSGW140 | celadussa ( C ) |

SNAPP run parameters: Mutation rates u and v were set to 1 and not sampled, coalescence rate was set to 10 and sampled. Non-polymorphic sites were excluded and log likelihood correction was used. Priors: alpha=11.75, beta=109.73, Kappa=1.0 and Lambda=10, all not sampled. MCMC chain length=1000000 Number of steps used during path sampling analysis=24.

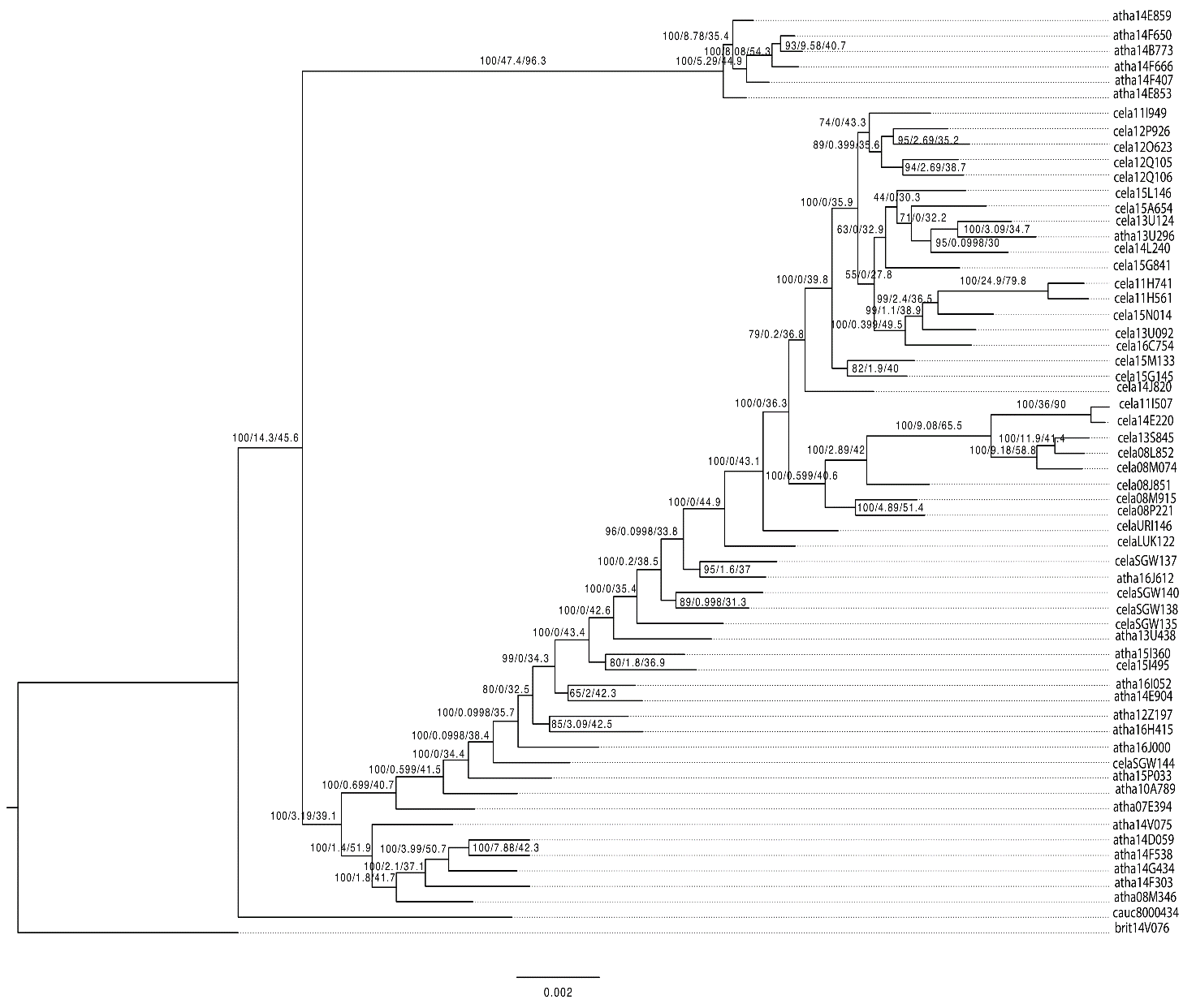

**Figure S1**: Species tree generated using the concatenation approach in IQTREE with bootstrap/gCF/sCF values labelled on branches

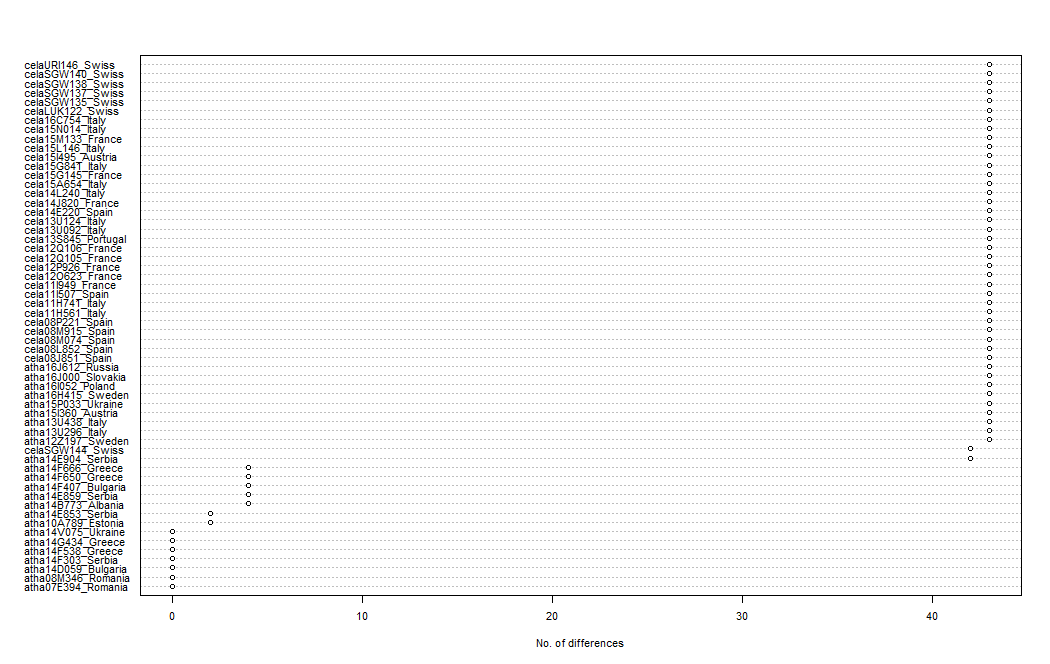

**Figure S2**: Dotplot of tip differences between TE60 and ddRAD trees obtained using the R function *tipdiff.*

*Indicating specimens from the contact zone*

In order to categorize the specimens belonging to the contact zone, we employed the biodecrypt approach, which identifies the areas of overlap among species based on convex hulls. We defined the contact zone as the intersection of the COI based hull distributions of *M. athalia* and *M. celadussa* and marked specimens belonging to the region of overlap as shown below.

**Table S2**: Specimens and their membership by location

| **ID** | **LAT** | **LON** | **Membership by location** |  |
| --- | --- | --- | --- | --- |
|  |  |  | 0 = contact zone or 50 km from it | 0 = contact zone or 100 km from it |
| atha14B773 | 40.592 | 20.596 | 2 | 2 |
| atha14E853 | 44.361 | 21.892 | 2 | 2 |
| atha14E859 | 44.361 | 21.892 | 2 | 2 |
| atha14F407 | 42.49 | 22.733 | 2 | 2 |
| atha14F650 | 40.205 | 21.064 | 2 | 2 |
| atha14F666 | 40.205 | 21.064 | 2 | 2 |
| atha07E394 | 44.812 | 25.397 | 2 | 2 |
| atha08M346 | 46.743 | 25.664 | 2 | 2 |
| atha10A789 | 58.107 | 26.918 | 2 | 2 |
| atha12Z197 | 59.612 | 18.485 | 2 | 2 |
| atha13U296 | 46.482 | 12.315 | 0 | 0 |
| atha13U438 | 46.589 | 12.853 | 0 | 0 |
| atha14D059 | 41.624 | 24.701 | 2 | 2 |
| atha14E904 | 44.156 | 19.693 | 2 | 2 |
| atha14F303 | 43.392 | 22.612 | 2 | 2 |
| atha14F538 | 41.371 | 23.633 | 2 | 2 |
| atha14G434 | 40.864 | 21.201 | 2 | 2 |
| atha14V075 | 49.817 | 35.75 | 2 | 2 |
| atha15I360 | 46.869 | 13.426 | 0 | 0 |
| atha15P033 | 50.31 | 29.11 | 2 | 2 |
| atha16H415 | 67.879 | 18.902 | 2 | 2 |
| atha16I052 | 52.819 | 14.235 | 2 | 2 |
| atha16J000 | 48.614 | 20.417 | 2 | 2 |
| atha16J612 | 54.386 | 22.368 | 2 | 2 |
| celaSGW144 | 47.208 | 9.388 | 0 | 0 |
| cela08J851 | 43.154 | -4.92 | 1 | 1 |
| cela08L852 | 40.878 | -3.848 | 1 | 1 |
| cela08M074 | 40.643 | -2.814 | 1 | 1 |
| cela08M915 | 42.765 | 0.712 | 1 | 1 |
| cela08P221 | 42.354 | 1.951 | 1 | 1 |
| cela11H561 | 38.095 | 13.249 | 1 | 1 |
| cela11H741 | 37.92 | 14.66 | 1 | 1 |
| cela11I507 | 37.083 | -3.51 | 1 | 1 |
| cela11I949 | 44.81 | 5.585 | 1 | 1 |
| cela12O623 | 44.201 | 7.074 | 1 | 1 |
| cela12P926 | 43.352 | 5.826 | 1 | 1 |
| cela12Q105 | 43.569 | 6.566 | 1 | 1 |
| cela12Q106 | 43.569 | 6.566 | 1 | 1 |
| cela13S845 | 40.389 | -7.534 | 1 | 1 |
| cela13U092 | 42.46 | 12.937 | 1 | 1 |
| cela13U124 | 46.024 | 12.28 | 1 | 0 |
| cela14E220 | 37.767 | -2.999 | 1 | 1 |
| cela14J820 | 43.782 | 2.727 | 1 | 1 |
| cela14L240 | 46.263 | 10.836 | 1 | 0 |
| cela15A654 | 45.328 | 9.509 | 1 | 1 |
| cela15G145 | 46.365 | 5.898 | 1 | 1 |
| cela15G841 | 46.294 | 8.28 | 1 | 0 |
| cela15I495 | 46.789 | 12.875 | 0 | 0 |
| cela15L146 | 46.424 | 10.464 | 1 | 0 |
| cela15M133 | 46.116 | 5.628 | 1 | 1 |
| cela15N014 | 41.449 | 15.112 | 1 | 1 |
| cela16C754 | 43.924 | 11.792 | 1 | 1 |
| celaURI146 | 46.907 | 8.52 | 0 | 0 |
| celaLUK122 | 47.016 | 8.253 | 0 | 0 |
| celaSGW135 | 47.208 | 9.388 | 0 | 0 |
| celaSGW137 | 47.208 | 9.388 | 0 | 0 |
| celaSGW138 | 47.208 | 9.388 | 0 | 0 |
| celaSGW140 | 47.208 | 9.388 | 0 | 0 |

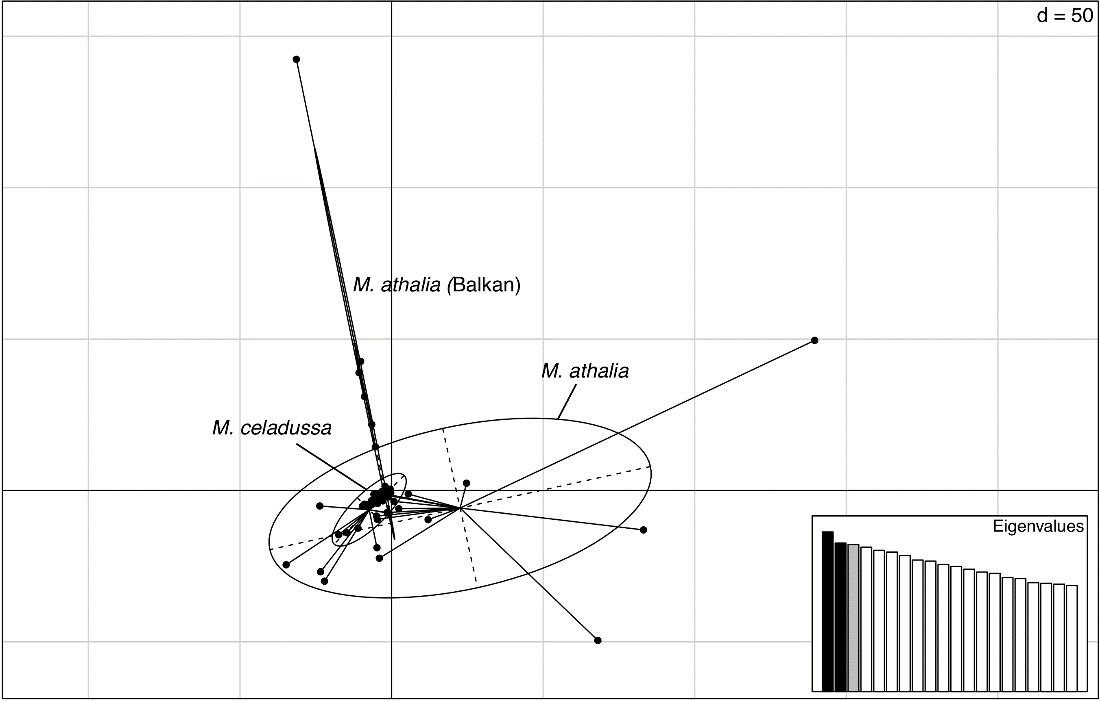

**Figure S3**: PCA plot for unlinked SNPs from ddRAD dataset

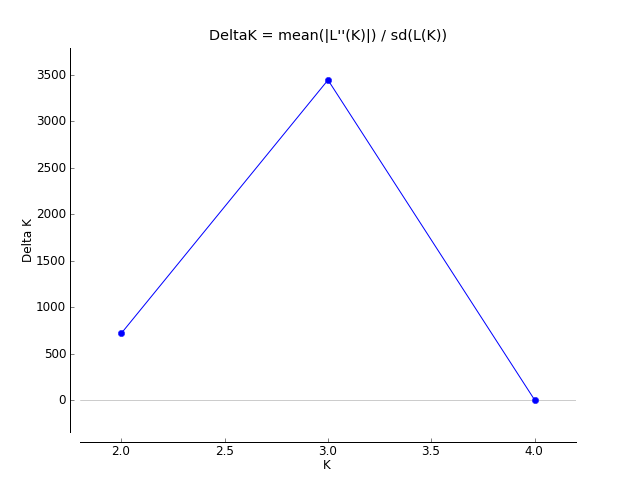

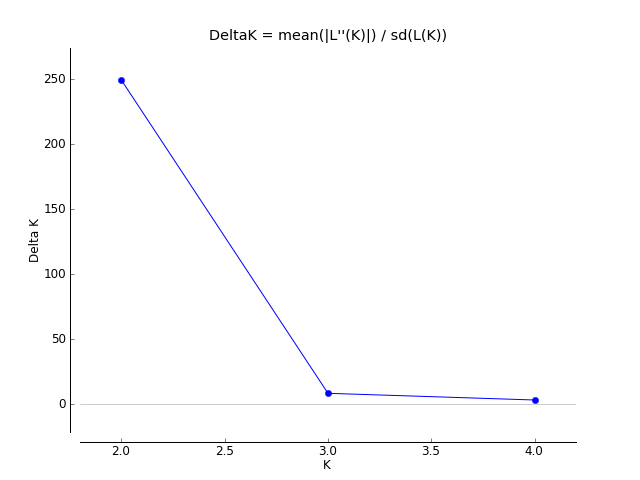

**Figure S4**: The ΔK plot for the ddRAD dataset showing optimum at K=2 (on right) and at K=3 (on left) for the target enrichment dataset.
